# RiboCollSensor: a sensitive real-time detector of ribosome collisions in mammalian cells based on split-NanoLuc complementation

**DOI:** 10.64898/2026.07.15.738639

**Authors:** José Alcalde, Iván Ventoso

## Abstract

Disruption of ribosome flux on translating mRNAs can result in ribosome collisions that activate key cellular responses. Despite growing interest in the field, current methods to detect ribosome collisions have limited sensitivity and are not suitable for use in living cells. Here, we describe a novel, reliable, and highly sensitive method based on split-nanoluciferase complementation to detect ribosome collisions in living cells. RiboCollSensor relies on the specific recruitment of EDF1-LgBiT to collided ribosomes near uS4-SmBiT, which generates luminescence due to the proximity of the partners in the ribosome. This biosensor showed unprecedented sensitivity, allowing detection of basal ribosome collisions in unstressed cells or under very low stress levels, enabling real-time analysis of collision kinetics. Thus, an increase in collisions could be detected within the first minute after translation disturbance, confirming the role of ribosomal flux as a rapid sensor of cell stress. Ribosome collisions rapidly disappeared after stress withdrawal, whereas under persistent stress, recovery was slower, taking up to two hours depending on the cell type. The ease and flexibility of this method, which requires only transient co-expression of sensor partners in target cells, make it applicable across many cell types to monitor the impact of internal and external cues on ribosome dynamics in real time.

## Introduction

Ribosomes not only synthesize proteins, but they also constitute a wide-ranging sensory network that detects stress and promotes downstream signaling to regulate proteostasis and cell fate. Several recent reports have confirmed the centrality of ribosome dynamics in stress sensing and mRNA fate, as well as their broad implications in cellular homeostasis (1–5). Ribosome flux on mRNAs can be disrupted by many stressors, including nutrient starvation— which reduces the pool of aminoacyl-tRNAs—UV irradiation, or low concentrations of elongation inhibitors (1, 6–9). Other sources of endogenous stress, including ROS production and reactive metabolic aldehydes, can damage mRNA by inducing oxidation of nucleobases (10). All these insults can promote ribosome stalling on translating mRNA, thereby increasing the probability of collision with trailing ribosomes. The resulting ribosome collisions are detected by several protein sensors recruited to collided ribosomes, including ZAKα, ubiquitin ligase ZNF593, EDF-1 (endothelial differentiation factor 1), and GCN1/GCN20 (1, 11–16). Moderate ribosome collision may induce translation initiation attenuation through GCN2-mediated eIF2α phosphorylation or by the recruitment of translation repressors GIGYP2/4EH1 to relieve ribosomal blockade (1, 15). However, strong collision can further trigger the ribotoxic stress response (RSR) through ZAKα-mediated activation of p38 and JNK MAPK, which induce cell cycle arrest and initiate apoptotis. The available structural models of collided disomes show the existence of an extensive interaction interface involving the solvent sides of both trailing and stalling 40S ribosomal subunits, the latter being markedly rotated relative to the former (13, 17). One of the collision sensors, EDF-1, is specifically recruited to collided ribosomes near the mRNA entry channel of the 40S subunit, where it contacts both uS3 (old RPS3) and h16 of 18S rRNA (15, 16, 18). EDF-1 binding can promote feedback inhibition of translation initiation at two levels: by recruiting translation repressors GIGYP2 and 4EH1 to collided ribosomes and by enhancing activation of the integrated stress response, which results in eIF2α phosphorylation (15, 18).

Current techniques to detect ribosome collisions involve the controlled digestion of polysomal mRNAs with RNase A to identify the accumulation of disomes or trisomes, either by sucrose gradient centrifugation or by RNAseq (disome profiling) (1, 19–21). However, these techniques are cumbersome and difficult to standardize as they generally require careful optimization of conditions to achieve controlled mRNA digestion. Moreover, polysome profiling by sucrose gradient centrifugation is not strictly a quantitative method and has limited sensitivity. Ubiquitination of eS10 and uS10 proteins by ZNF598 has also been used as a readout of ribosome collision; however, this modification can also occur on stalled ribosomes—not necessarily colliding—as part of the ribosome-associated quality control (RQC) (13, 14, 22). A serious limitation of all these procedures is that they are not amenable to real-time detection of ribosome collisions in living cells, which may occur soon after a disturbance in ribosomal flux.

Among the reporter genes used as sensors, nanoluciferase (NanoLuc) showed the highest activity and sensitivity (23). In 2016, a split-NanoLuc complementation assay was developed to detect protein–protein interactions with unprecedented sensitivity (NanoBiT) (23–25). Later, long-lived variants of the NanoLuc substrate, furimazine, were developed to allow sustained luminescence measurements in living cells. Here, we used this technology to design a simple, sensitive method to detect ribosome collisions in living cells.

## Materials and Methods

### Plasmid construction

To construct sensor-expressing plasmids, we first subcloned SmBiT and LgBiT DNA fragments encoding the small and large subunits of NanoLuc, respectively, into a plasmid under the CMV promoter (pSmBiT and pLgBiT). For this, the *Hind*III-*Xba*I EGFP fragment of the pEGFP-N1 plasmid (Clontech) was replaced by the corresponding DNA fragments of Sm/LgBiT from pBIT1.1 and pBIT2.1 plasmids (NanoBiT, Promega), respectively, using the same enzymes. Then, cDNA sequences of human *Rack1*, *uS5* (old RPS2), *uS3* (old RPS3), *uS4* (old RPS9), and *EDF1* genes were obtained by retrotranscription of total RNA from HEK293T or HeLa cells using SuperScript IV RT (Invitrogen), followed by standard PCR amplification (Supreme NYZTaq II, NZYtech). The primers used are listed in the supplementary information. All constructs were verified by sequencing.

### Cell culture and transfection

HEK293T, HeLa, and MEFs cells were grown in DMEM supplemented with antibiotics and 10% fetal calf serum under standard conditions. Cells were authenticated by microscopic examination, and they were free of mycoplasma contamination. Cells growing in 24-well plates at 70–80% confluency were co-transfected with a combination of 0.25 µg of each pSm-X and pLg-Y plasmids and 1 µL of TurboFect (Thermo Fisher Scientific) or 1.5 µL of Lipofectamine 2000 (Invitrogen). Lipofectamine 2000 was chosen for the transfection of MEFs. We also tested Lipotransfectin (NiborLab) with similar results. Twenty-four hours later, the cells were trypsinized and split into 48-well or 96-well plates for analysis the following day.

### Cell treatment with stressors

Anisomycin and emetine (both from SIGMA) were dissolved in DMSO and used at final concentrations of 0.001–3 and 0.5–5 µM, respectively. For real-time measurements, the injectors of the GloMax reader were primed with Opti-MEM containing drugs at a 10-fold final concentration, and 10-15 µL was injected per well. For UVC treatment, cell plates were irradiated at 10 cm from the 257 nm lamps in a UV crosslinker (BioRad), which was previously calibrated using a radiometer (UVX, UVP). At the indicated distance, the cell plates received a radiation intensity equivalent to 250 μW/cm^2^. Cells were irradiated for 7 to 90s, resulting in cumulative intensities from 1.75 to 22 mJ/cm^2^. For amino acid deprivation, we prepared DMEM without leucine or arginine and supplemented with dialyzed fetal bovine serum (FBS) as described previously. Cells were washed twice with medium lacking the indicated amino acid and incubated for up to 10 hours.

### Measurement of NanoLuc activity in cell extracts

Forty-eight hours post-transfection, cells were washed with phosphate-buffered saline (PBS) and lysed in 100 µL (for L-24 and L-48 wells) or 50 µL (for L-96 wells) cold polysome lysis buffer (Tris-HCl 25 mM pH 7.5, 0.1M KCl, 10mM Mg(CH_3_COO)_2_, 0.5% Triton X-100, 2mM DTT and 250 µM anysomycin) for 15 min on ice. Luciferase activity was measured directly using 20 µL of cell extract and 20 µL of reaction buffer (polysome buffer without Triton-X100 and anisomycin) containing furimazine substrate (Nano-Glo, Promega). Depending on the number of samples to be measured, we used a Berthold luminometer or a GloMax Discover plate reader (Promega) with an integration time of 0.3 s.

### Real-time measurement of NanoLuc in living cells

To analyze NanoLuc activity *in vivo*, transfected cells were plated on CellCarrier-96 Ultra microplates (PerkinElmer) the day before analysis. The medium was replaced with 150 μL of Opti-MEM medium containing 2% FBS and the appropriate dilution of Nano-Glo Vivazine substrate (Promega). The cells were incubated for 2 hours at 37°C before analysis to allow substrate stabilization. Then, the plate was transferred to the GloMax Discover reader previously set to 37°C, and NanoLuc activity was recorded at 30 s, 1 min, and 2 min intervals.

### Detection of chimeric proteins by western-blot

To detect the expression of tagged proteins, cells were lysed in sample buffer 48 hours post-transfection, and analyzed by 12% sodium dodecyl sulfate– polyacrylamide gel electrophoresis (SDS-PAGE) followed by wet transfer to a PVDF membrane (Immobilon, Merck) as described previously (26). Primary antibodies used were: anti-EDF1 (St Johńs Lab # STJ73046 and Abcam #174651), anti-RPS9 (Santa Cruz Biotech # 390614), anti-eIF2 (Cell Signaling #9722), anti-eIF2-P (Cell Signaling # 9721S), anti-p38 (Cell signaling #9218) and anti-p38-P (Cell Signaling #9212S). Primary antibodies were incubated overnight at 4°C in BLOTTO buffer (TBS 0.05% Tween-20+ 5% milk), washed three times in BLOTTO without milk, and incubated with secondary antibodies conjugated to POD (Promega) for 1 hour at room temperature. After three final washes, chemiluminescence was detected using Luminata Crescendo (Millipore) in an ImageQuant apparatus (GE).

### Polysome profile analysis

One P100 plate of subconfluent HEK293T cells was transfected with 2 µg of each plasmid expressing EDF1-Lg or RPS9-Sm, and 24 hours later, the cells were split between two P100 plates and incubated for one more day. Forty-eight hours post-transfection, one plate was treated with 0.5 µM anisomycin for 20 min at 37°C, washed with cold PBS, and lysed in 1 mL of cold polysome buffer supplemented with RNase inhibitor (NYZTech) for 15 min on ice. Cell lysates were clarified by low-speed centrifugation and loaded on a 10–40% sucrose gradient prepared in polysome buffer without Triton X-100. Gradients were centrifuged at 35,000 rpm for 3 hours at 4°C in a Beckman SW40.1 rotor and fractionated from the top using an ISCO fractionator coupled to a UV recorder. A 50 µL sample of each fraction was used to measure NanoLuc activity as described above. For protein analysis, a 1mL fraction was precipitated in 15% trichloroacetic acid (TCA) overnight at -20°C, washed twice with acetone, and dissolved in sample buffer for western blot analysis as described above.

### Structural modeling

Using the recently reported structural model of ZAK disome purified from ANS-treated cells (PDB 9RPV), we modeled EDF1-Lg and uS4-Sm into the colliding 40S ribosomal subunit using the Matchmaker function of Chimera. Models of EDF1-Lg and RPS9-Sm were generated using AlphaFold.

### Statistical analysis

Graph construction and statistical analyses were performed using GraphPad Prism 6. Two-tailed *t*-test using at least three technical replicates per sample or three biological replicates per group. Significance was scored as follows: *P* < .0001, \*\*\**P* < .001, \*\**P* < .05, \**P* > .05, ns = not significant.

## Results

### Identification of ribosome collision readers

Among the ribosomal proteins and recruited factors that are located at or near the collision interface of disomes (PDB 9RPV), we selected Rack1, uS5 (old RPS2), uS3 (old RPS3), uS4 (old RPS9), and EDF1 to find partners that could generate NanoLuc complementation (Figure 1a). For this, we tagged selected proteins with LgBiT or SmBiT fragments at the N- or C-terminus, followed by pairwise coexpression in HEK293T cells by transfection. We set two criteria to identify those NanoLuc complementations that can occur specifically in colliding ribosomes. First, a strong and reproducible increase in NanoLuc should be detected after the addition of known inducers of ribosome collision (e.g., low concentration of anisomycin, ANS). Second, since a low basal level of ribosome collisions is expected in untreated cells according to previous reports (19, 27), those partners that faithfully read out ribosome collisions should render a low but detectable basal activity. Therefore, this basal activity should be sensitive to polysome breakdown by high doses of puromycin. Coexpression of some partners resulted in strong NanoLuc complementation in control cells due to their proximity to 40S ribosomal subunits, as occurred for uS5-Lg and uS3-Sm (Figure 1b). Interestingly, the combinations EDF1-Lg and uS4-Sm, and to a lesser extent, EDF1-Sm and uS4-Lg, fulfilled our criteria as ribosome collision readers. Thus, a strong increase in NanoLuc activity was specifically detected 15 minutes after the addition of 0.3 µM ANS, together with detectable NanoLuc activity in control cells, which was reduced by puromycin treatment (Figure 1b). The EDF1-Lg and uS5-Sm combination also resulted in an increase in luciferase activity upon ANS treatment, although to a lesser extent than the EDF1-Lg + uS4-Sm combination. Since EDF1 has been detected bound to the collided, but not to the stalled ribosome in structural models of disomes, the NanoLuc complementation observed can easily be explained by the recruitment of EDF1-Lg near uS4-Sm within the collided ribosome. Indeed, the C-terminal ends of EDF1 and uS4 are arranged towards the same side in the collided 40S subunit and separated by a distance (50–60 Å), compatible with NanoLuc complementation (Figure 1c) (23).

**Figure 1.**
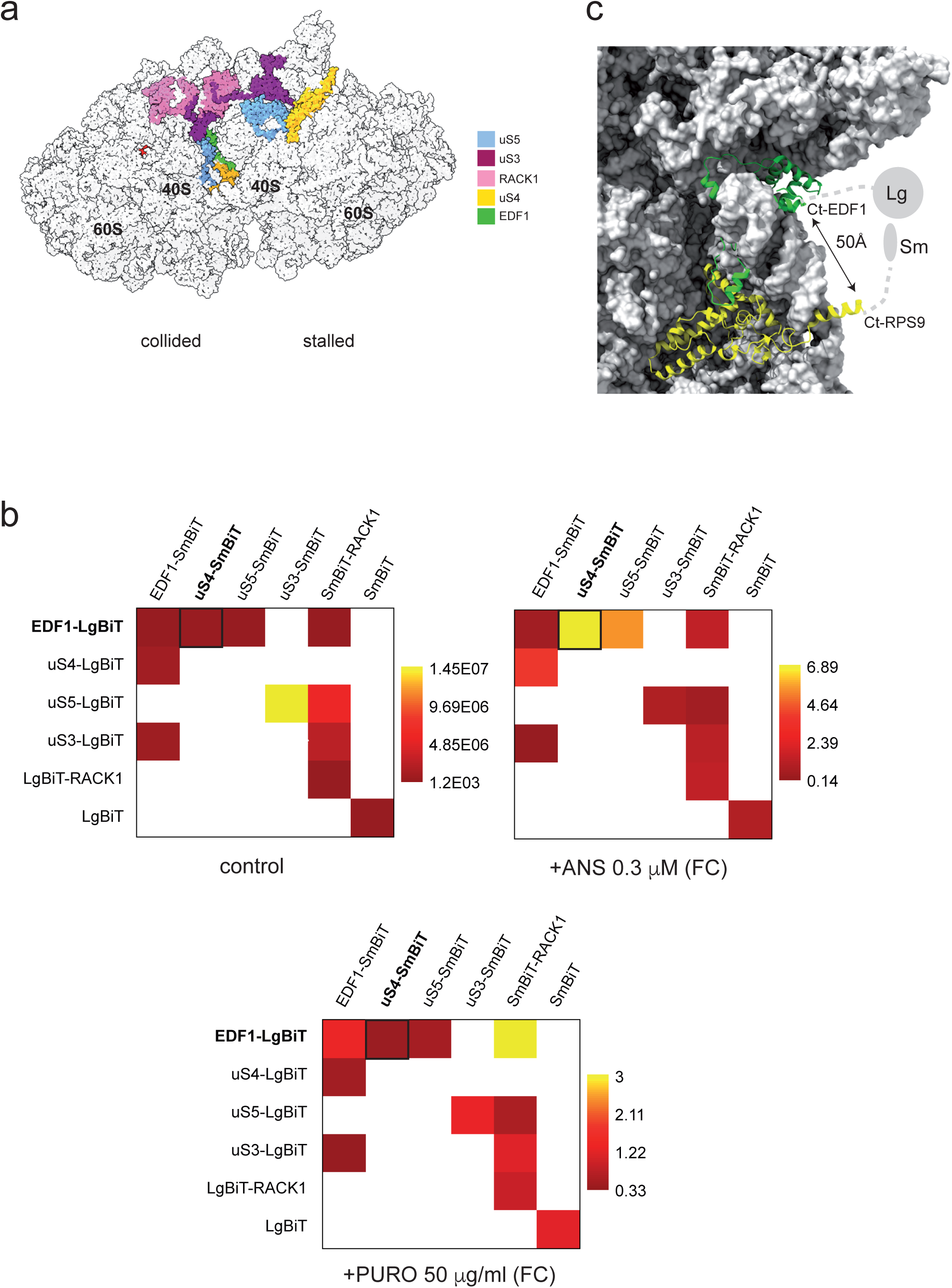
(a) Model of a colliding disome (PDB 9RPV) showing some of the ribosomal proteins and recruited factors (EDF1) located at or near the collision interface. Those analyzed here are colored in the legend. (b) Pairwise analysis of NanoLuc complementation. The indicated pairs of protein-expressing plasmids were transfected into HEK293T cells and analyzed 48 hours later. Empty cells in the matrix correspond to untested pairs. Cells were untreated (control), treated with 0.3 μM anisomycin (ANS), or with 50 μg/ml Puromycin (PUR) for 20 min. For ANS- and PURO-treated cells, data are expressed as fold change (FC) with respect to the control. (c) Close view of EDF1 bound to a collided 40S subunit showing the proximity to uS4. The estimated distance between the projected C-terminal ends of EDF1 and uS4 is indicated.

### RiboCollSensor performance and reliability

Next, experiments were undertaken to characterize the sensor based on the combination of EDF1-Lg and uS4-Sm (Figure 2a). First, we quantified the expression of EDF1-Lg and uS4-Sm proteins in co-transfected cells by western blot. In HEK293T cells, uS4-Sm and EDF1-Lg chimeras accumulated two or three times more than the endogenous counterparts, although for EDF1 the comparison was not reliable since the endogenous form of this protein (17 kDa) was not efficiently detected by either the two different antibodies tested (Figure 2b). To test the sensor’s performance and reproducibility with known inducers of ribosome collisions, we also included UV radiation in addition to a low dose of ANS (1, 8). Notably, both treatments induced a robust increase in NanoLuc activity in both HEK293T cells and mouse embryonic fibroblasts (MEFs), ranging from 10- to 100-fold depending on the cell type used (Figure 2c). Among the collision inducers tested, ANS was the strongest. We confirmed that these stress treatments induced an RSR through p38 activation and eIF2α phosphorylation, as previously described (Figure 2d).

**Figure 2.**
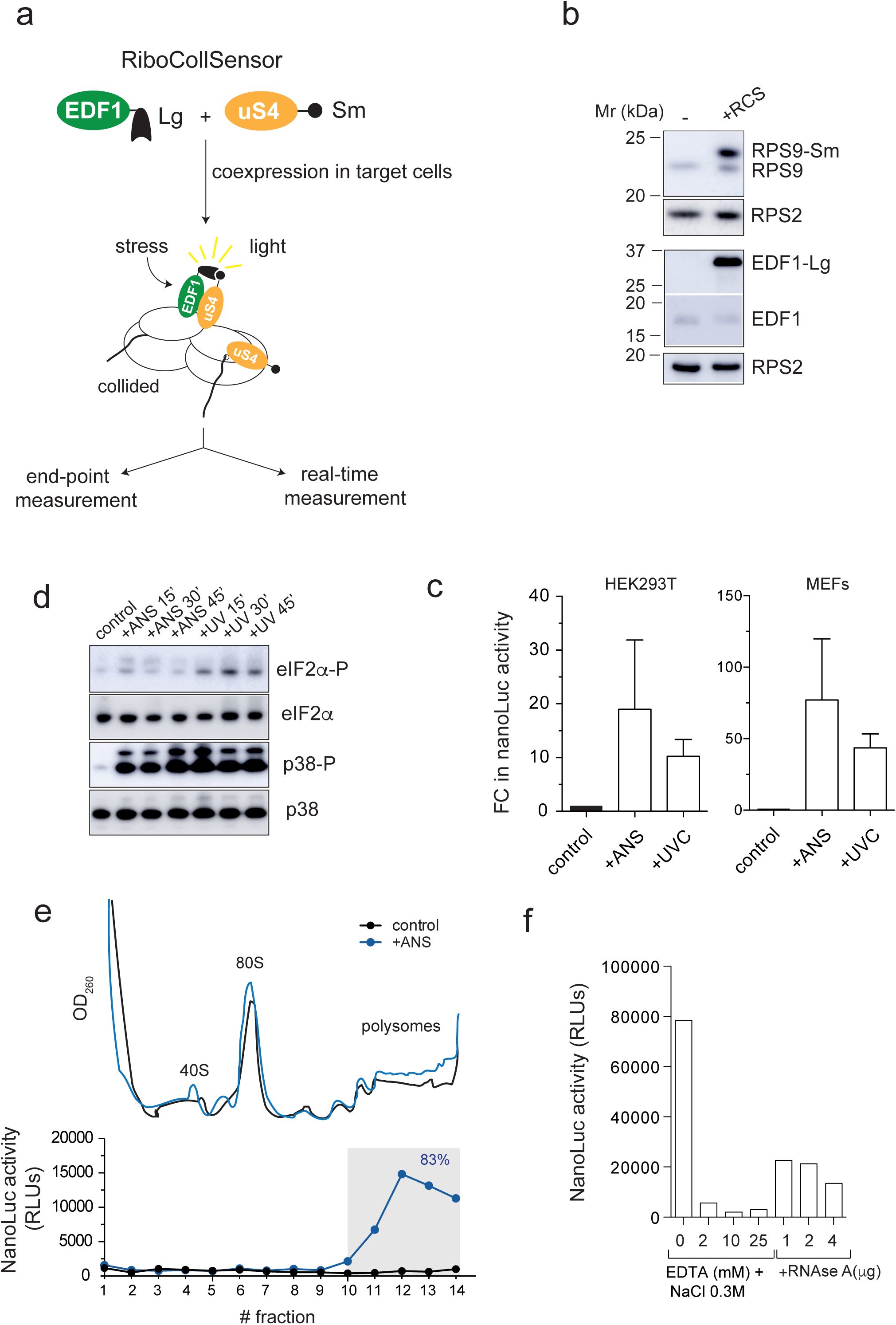
Testing RiboCollSensor reliability. (a) Schematic protocol to measure ribosome collisions based on co-expression of EDF1-Lg and uS4-Sm in target cells. (b) Accumulation of uS4-Sm and EDF1-Lg proteins in transfected HEK293T cells analyzed by western blot. To detect endogenous EDF1, the corresponding part of the blot was exposed longer (lower panel). (c) Response of RiboCollSensor to ANS (0.3 µM) and UVC radiation (30mJ/cm^2^) in HEK293T cells and MEFs. Measurements were taken after 15 min of treatment. Data are the mean fold change (FC) ± SD from at least five independent experiments. (d) Western-blot analysis of p38 and eIF2α phosphorylation in response to ANS and UVC. (e) Polysome profiles of control (black) and ANS-treated cells (blue) analyzed by ultracentrifugation in sucrose gradients. Peaks corresponding to ribosomal subunits, monosomes, and polysomes are indicated. NanoLuc activity detected in 25 μL-sample of the resulting fractions is also shown (bottom). (f) Sensitivity of NanoLuc activity to polysome breakdown. Samples of pooled polysome fractions (35 μL, about 2 μg total RNA) from ANS-treated cells were incubated with EDTA/NaCl 0.3M or increasing amounts of RNase A for 15 min at room temperature, and the resulting NanoLuc activities were measured.

To confirm that RiboCollSensor reads out true ribosome collisions, we analyzed NanoLuc activity in fractions of polysome sucrose gradients in both control and ANS-treated HEK293T cells. As expected, NanoLuc activity was detected exclusively in polysomes but not in 80S or in free ribosomal subunits, since collisions must occur on translating polysomal mRNA (Figure 2e). Similar results were found in MEFs (Figure S1). To further confirm this point, we disrupted polysomes from ANS-treated cells with a combination of EDTA and 0.3M NaCl, resulting in a complete loss of NanoLuc activity. Accordingly, parallel treatment with increasing amounts of RNase A only partially reduced NanoLuc activity, since collided ribosomes are resistant to mild RNase A treatment (Figure 2f).

However, no NanoLuc activation was detected by leucine or arginine deprivation at any of the time points analyzed in either HEK293T cells or MEFs (Figure S2). Since these treatments induce rapid eIF2α phosphorylation due to autonomous, non-collision–dependent activation of GCN2—which could reduce translation initiation and collisions—we repeated the experiment in eIF2α S51A knock-in (KI) MEFs (28). A low but significant increase in NanoLuc activity was detected in these cells, indicating that RiboCollSensor can also detect ribosome collisions caused by an aminoacyl-tRNA shortage under some circumstances (Figure S2).

### RiboCollSensor sensitivity

To test the sensitivity of RiboCollSensor, we titrated the number of sensor-expressing plasmids and stress dose to find the lower limit of response. Significant NanoLuc activation by ANS treatment was achieved using as little as 4 ng of each sensor plasmid in 10^5^ cells, despite the very low basal activity detected under this condition (Figure 3a). Notably, the sensor could detect a robust increase in NanoLuc activity using a very low dose of ANS (12 nM) or UVC (2 mJ/cm^2^), more than 20-fold lower than those typically used to detect ribosome collisions by other methods (Figures 3b and c). These results demonstrated the exquisite sensitivity of the sensor.

**Figure 3.**
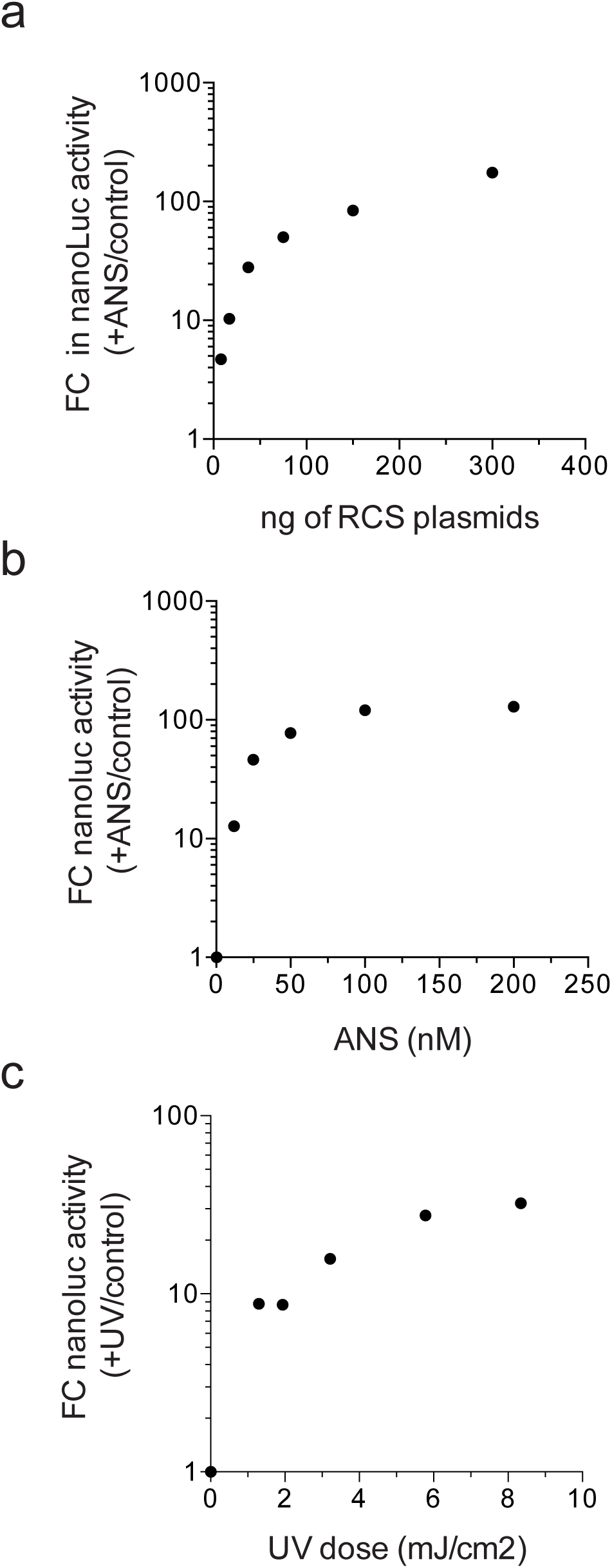
(a) Influence of the number of sensor-expressing plasmids on RiboCollSensor response. MEFs were transfected with the indicated amount of RCS plasmids (at a 1:1 ratio) and treated or not with ANS 0.3 μM for 15 min. RiboCollSensor response to increasing concentrations of ANS (b) and UVC doses (c). In all cases, the resulting FC in NanoLuc activity with respect to unstressed cells is shown.

### Analysis of ribosome collision dynamics

Next, we closely tracked the appearance and disappearance of ribosome collisions in real-time experiments. Maximal activation by ANS was reached at 20-30 min in MEFs and 15 min in HEK293T cells, followed by a slower decline that lasted 2–3 hours for MEFs (Figure 4a). However, stressor withdrawal after 15 min allowed the rapid disappearance of ribosome collisions following a similar but inverse kinetics. To gain further insights into this process, we analyzed the appearance of ribosome collisions in greater detail. We consistently found an increase in NanoLuc activity within the first minute after ANS injection (Figure 4b). Assuming that ANS uptake into the cells is a non-instantaneous process, our data suggest that cells can begin to detect ribosome collisions after seconds of exposure to stress.

**Figure 4.**
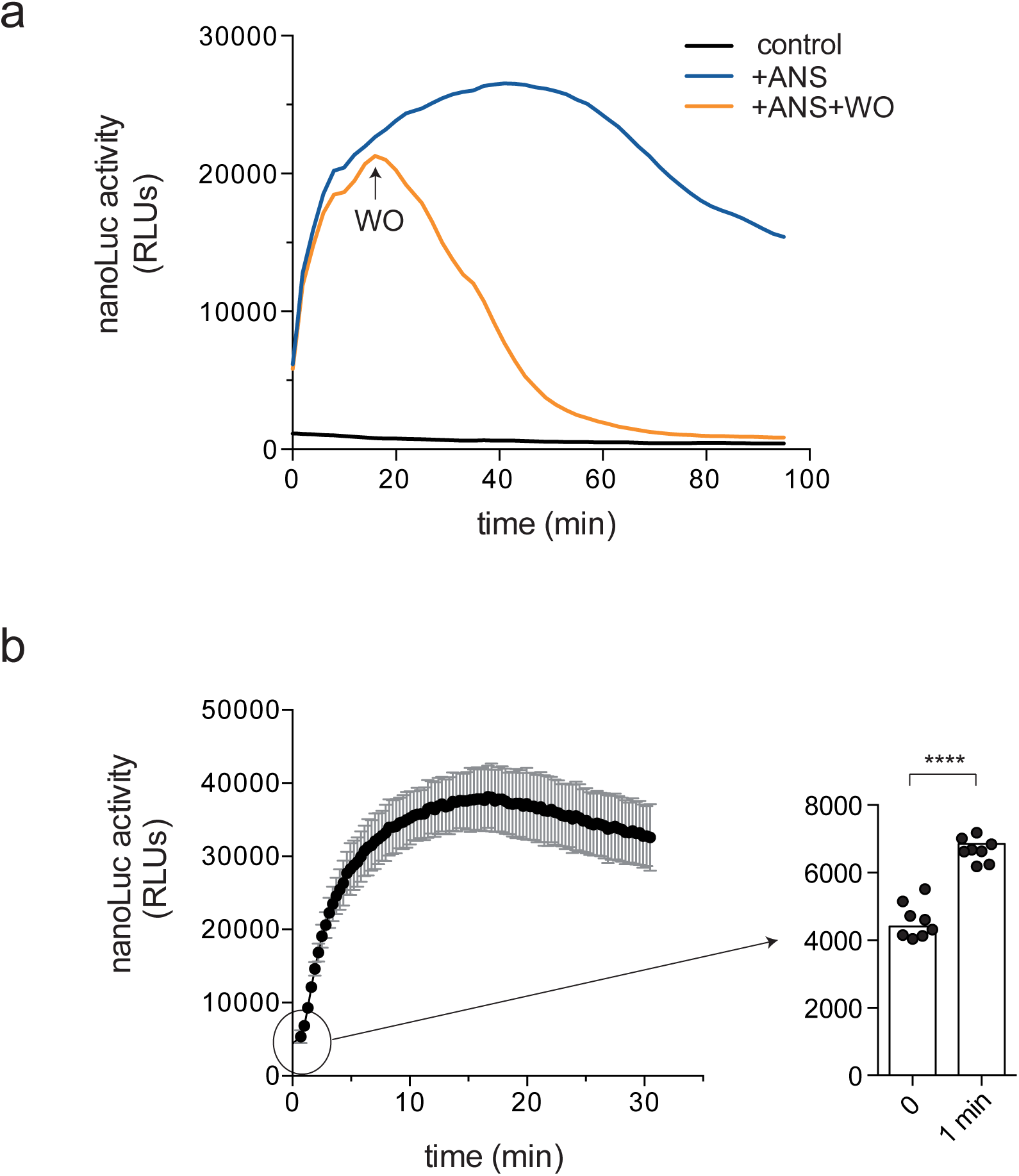
(a) Ribosome collision dynamics analyzed by real-time experiments. MEFs were transfected with plasmids encoding sensors, and 48 hours later, collisions were induced with ANS 0.3 μM (blue and orange lines). NanoLuc activity was measured every 2 minutes as described in the methods. Where indicated (WO, orange line), ANS was washed out after 15 min of treatment, and measurement was continued. (b) Short-time analysis of collision kinetics. HEK293T cells expressing sensors were treated with 0.3 μM ANS, and NanoLuc activity was recorded every 30 s. Measurements were taken in quadruplicate, and the resulting means and standard deviations (SDs) were used to generate the curve. A close-up measurement at 1 min after ANS injection is also shown, including a t-test comparing control and ANS-treated samples.

### Detection of basal ribosome collisions in unstressed cells

A basal level of ribosome collisions is expected in unstressed cells as a consequence of active translation, as reported previously (19, 20). To roughly quantify this, we first measured NanoLuc activity in cells treated with high puromycin (50 µg/ml), which runs off polysomes, thus reducing ribosome collisions to 0 (Figure 5a). The resulting residual activity was considered noise of the NanoBit system. Notably, a 2- to 3-fold reduction in NanoLuc activity relative to control cells was observed, indicating the existence of basal ribosome collisions. Likewise, treatment with high concentrations of translation inhibitors, such as harringtonine (HT), reduced NanoLuc activity to a similar extent, as expected, since initiation blockade relieves ribosome collisions on translating mRNAs (Figure 5b). However, since RiboCollSensor can not capture those collisions involving ribosomes lacking tagged EDF1/uS4, an absolute quantification of basal ribosome collisions was not possible. Therefore, we conducted a relative estimation of basal collisions with respect to maximal collisions found using low doses of ANS. Thus, we estimated that basal ribosome collisions represented about 7% and 1% of maximal collisions observed upon ANS treatment in HEK293T cells and MEFs, respectively.

**Figure 5.**
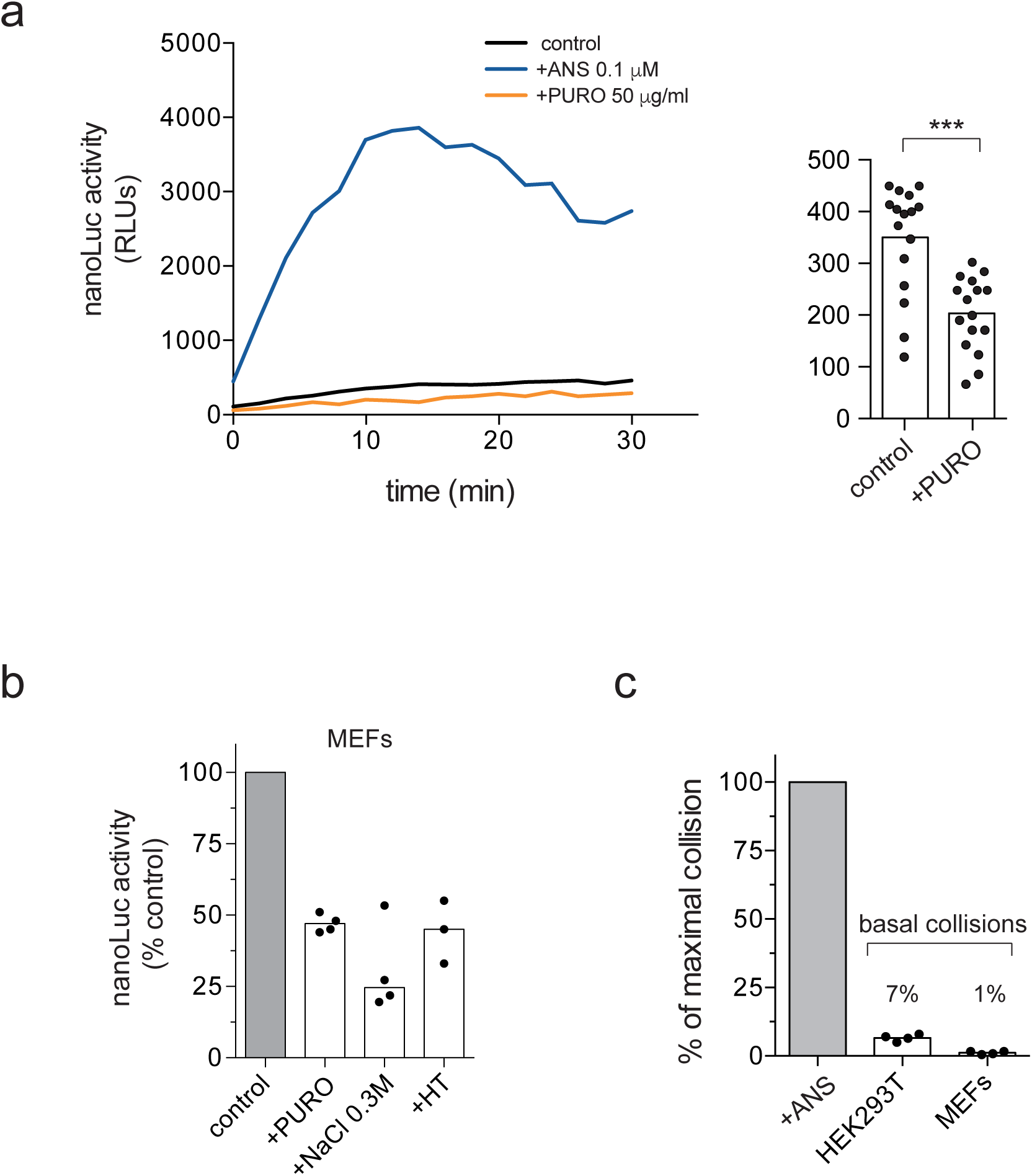
(a) Measurement of basal ribosome collisions. HEK293T cells expressing sensors were treated with 0.1 μM ANS or 50 μg/mL PURO, and NanoLuc activity was recorded every 2 min. The resulting quadruplicate measurements are presented as means, including a t-test analysis comparing control and PURO-treated cells. (b) Effect of high-dose PURO, NaCl 0.3M, and harringtonine (30μg/ml) treatments on NanoLuc activity in MEFs. (c) High-dose PURO treatment allowed relative quantification of basal ribosome collisions. The activity in control cells was subtracted from that in PURO-treated cells, and the resulting activity was normalized to the maximal activity found in ANS-treated cells across four independent experiments.

## Discussion

The reliability of the sensor presented here for detecting true ribosome collisions is supported by three main results. First, a strong increase in NanoLuc was consistently observed upon treatment of different cell types with known ribosome-collision inducers. Second, NanoLuc complementation was only detected in polysomes, not in 80S monosomes, as expected, since ribosome collisions must occur on translating mRNAs with two or more bound ribosomes. Third, NanoLuc activity detected in polysomal fractions was highly sensitive to dissociative treatments (EDTA/NaCl) but partially resistant to RNase A treatment. Furthermore, since RiboCollSensor depends on the specific recruitment of EDF1-Lg to collided ribosomes, ectopic NanoLuc complementation arising from 80S proximity within densely packed polysomes is expected to contribute minimally, if at all, to sensor activity.

One of the main features of the sensor is its high sensitivity in detecting ribosome collisions at very low stressor doses and over very short time periods. Even though RiboCollSensor could not capture all ribosome collisions that may occur in the cell, this technology was sufficiently sensitive to detect basal ribosome collisions in unstressed cells, which represent only a small fraction (1– 7%) of the maximal activity found in stressed cells. This aligns with genome-wide analysis in several species showing that a significant fraction of 80S bound to mRNAs exhibited the typical disome fingerprint of colliding ribosomes, mostly associated with an inefficient stop codon context in some mRNAs and the presence of certain amino acid motifs in nascent polypeptides (19, 27).

Real-time recordings using the sensor revealed ribosome collisions as a highly dynamic process that can quickly detect translation disturbances. The existence of a rapid response to UV or to low concentrations of elongation inhibitors had been anticipated in several reports (1, 8); however, RiboCollSensor now shows that this response may be even faster than previously thought. Similarly, ribosome collisions vanished upon stress withdrawal, whereas in the continued presence of stressors, cells could still significantly resolve ribosome collisions within a few hours, particularly in HEK293T cells. All these findings place ribosomal dynamics not only at the center but also at the forefront of cellular stress response. However, RiboCollSensor was unable to detect ribosome collisions after amino acid deprivation in HEK293T and MEFs, whereas only a modest increase in NanoLuc activity was detected in eIF2α S51A KI MEFs. This agrees with previous reports showing that amino acid deprivation alone was not an effective inducer of ribosome collisions (1, 3, 21). This was probably due to the collision-independent branch of GCN2 activation triggered by the accumulation of uncharged tRNAs, which can also promote a rapid blockade of translation initiation (29). Another possibility is that EDF1 may not be involved in sensing ribosome collisions due to amino acid deprivation, since, to our knowledge, this association has not been reported.

Since we found differential responses of MEFs and HEK293T cells to ribosome collision inducers, the flexibility of this technology will allow a systematic analysis of how cell type or metabolic status influences ribosome collision dynamics and stress responses. The low number of sensor plasmids required for detection would enable the use of RiBoCollSensor across a wide variety of cell types, including primary cells, such as neurons and lymphocytes, that are difficult to transfect. Finally, the use of this technology could allow for a systematic screening of new physical or chemical disturbances that might have a primary impact on ribosome flux, including the effects of certain pathological proteins involved in neurodegenerative disorders (9, 30).

## Acknowledgments

We thank the staff of our laboratory for their suggestions and for sharing ideas.

## Author contributions

I.V. conceived the study, directed the research, performed most experiments, and wrote the manuscript. J.A. cultured MEF cells and performed all western blots.

