## supplemental information for "RiboCollSensor: a sensitive real-time detector of ribosome collisions in mammalian cells based on split-NanoLuc complementation"

**Supplemental data**

Figure S1
Figure S2
Oligonucleotide sequences
Sequence of chimeric proteins

Figure S1

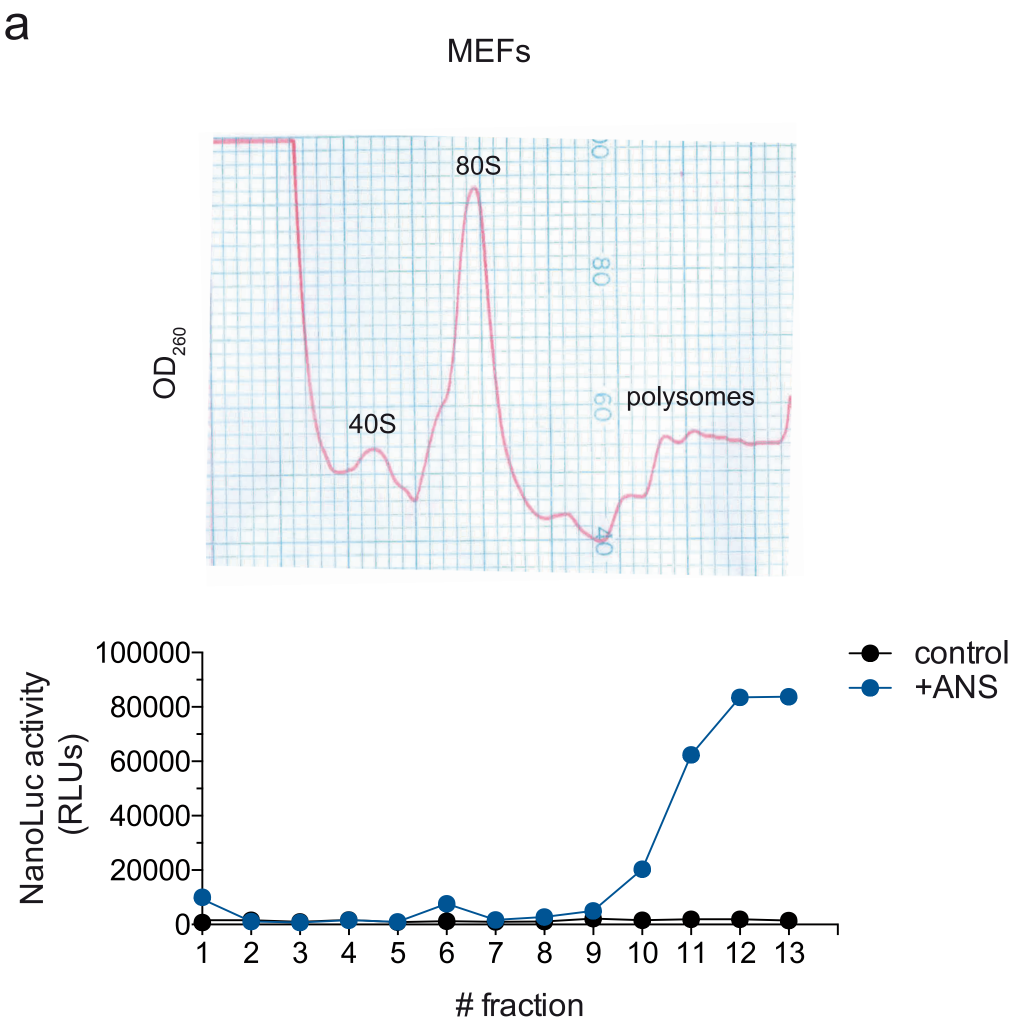


Figure S1. Polysome profiles of control (black) and ANS-treated MEFs (blue) analyzed by ultracentrifugation in sucrose gradients. Peaks corresponding to ribosomal subunits, monosomes, and polysomes are indicated. NanoLuc activity detected in the resulting fractions is also shown (bottom).

Figure S2

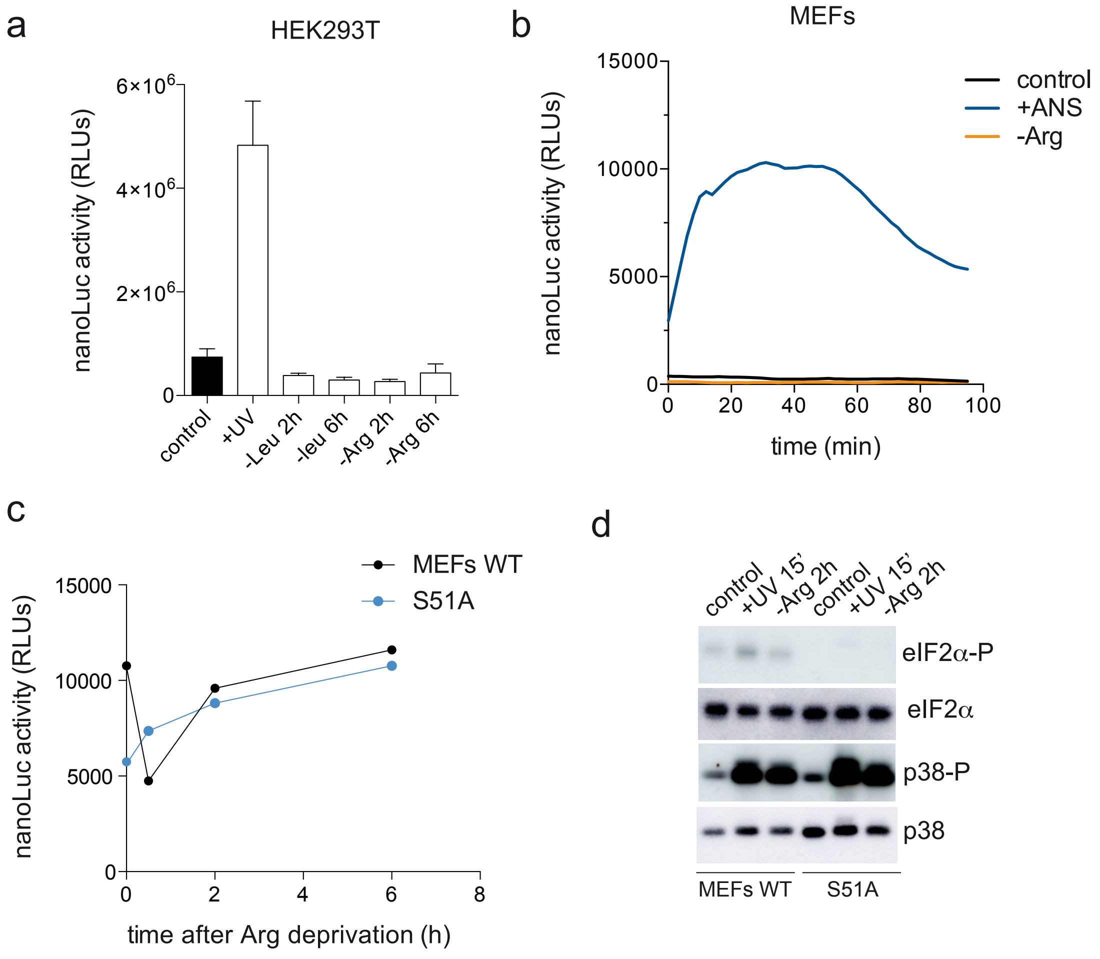


Figure S2. RiboCollSensor response to amino acid deprivation. (a) NanoLuc activity after 2 or 6 hours of leucine or arginine deprivation in HEK293T cells. UVC radiation was included as a positive control of ribosome collision. Data are the mean of triplicate experiments. (b) Real-time analysis of NanoLuc activity in MEFs deprived of arginine. Treatment with low-dose ANS was included as a positive control. (c) NanoLuc activity by arginine deprivation in MEFs WT or KI S51A. (d) Western-blot analysis of p38 and eIF2α phosphorylation in response to the indicated treatments.

### Oligonucleotide sequences

| | |
|---|---|
| fw-hEDF1 | CACACGAATTCAGACAAGCTTACC**ATGGCCGAGAGCGACTGGGACACGGTGACGG** |
| rev-hEDF1 | GGTGTGAATTCTGTCTAGA**TTTCGCCCTAGGCCCCTTCTCGATGG** |
| fw-hRPS9 | CACAACAAGCTTACC**ATGGCCCGGAGCTGGGTTTGTCGC** |
| rev-hRPS9 | GCCGCTCGAATT**CCCAGCCCCACCCTGGCCCTTCTTGGCATTC** |
| fw-hRPS2 | CACAACGAATTCCAAGCTTACC**ATGGGGAACCGCGGTGGCTTCCGCGGAGG** |
| rev-hRPS2 | ACACCACGAATTCCC**GGTGTGGGTCTTGACGAGGTGGTCAGTGA** |
| fw-hRPS3 | CACAACGAATTCCAAGCTTACC**ATGGCAGTGCAAATATCCAAGAAGAGGAAGTTTGTCG** |
| rev-hRPS3 | GTTGTCTAGATTAGCTCGAGTT**TGCTGTGGGGACTGGCTGGGGCATGGCAGGCGGCTC** |
| fw-RACK1 | CACACGAGCTCAAAAGCTTACC**ATGACTGAGCAGATGACCCTTCGTGGCACCCTC** |
| rev-RACK1 | GTTGTCTAGATTA**GCGTGTGCCAATGGTCACCTGCCACACTCGCACCAG** |

### Sequence of chimeric proteins. Human protein sequences (underlined) tagged with Sm or Lg (shaded in yellow) are shown.

EDF1-LgBiT

MAESDWDTVTVLRKKGPTAAQAKSKQAILAAQRRGEDVETSKKWAAGQNKQHSITKNTA
KLDRETEELHHDRVTLEVGKVIQQGRQSKGLTQKDLATKINEKPQVIADYESGRAIPNN
QVLGKIERAIGLKLRGKDIGKPIEKGPRAKSRQNSGSSGGGGSGGGGSSGVFTLEDFVG
DWEQTAAYNLDQVLEQGGVSSLLQNLAVSVTPIQRIVRSGENALKIDIHVIIPYEGLSA
DQMAQIEEVFKVVYPVDDHHFKVILPYGTLVIDGVTPNMLNYFGRPYEGIAVFDGKKIT
VTGTLWNGNKIIDERLITPDGSMLFRVTINS

EDF1-SmBiT

MAESDWDTVTVLRKKGPTAAQAKSKQAILAAQRRGEDVETSKKWAAGQNKQHSITKNTA
KLDRETEELHHDRVTLEVGKVIQQGRQSKGLTQKDLATKINEKPQVIADYESGRAIPNN
QVLGKIERAIGLKLRGKDIGKPIEKGPRAKSRQNSGSSGGGGSGGGGSSGVTGYRLFEE
IL

RPS9-LgBiT

MARSWVCRKTYVTPRRPFEKSRLDQELKLIGEYGLRNKREVWRVKFTLAKIRKAARELL
TLDKKDPRRLFEGNALLRRLVRIGVLDEGKMKLDYILGLKIEDFLERRLQTQVFKLGLA
KSIHHARVLIRQRHIRVRKQVVNIPSFIVRLDSQKHIDFSLRSPYGGGRPGRVKRKNAK
KGQGGAGNSGSSGGGGSGGGGSSGVFTLEDFVGDWEQTAAYNLDQVLEQGGVSSLLQNL
AVSVTPIQRIVRSGENALKIDIHVIIPYEGLSADQMAQIEEVFKVVYPVDDHHFKVILP
YGTLVIDGVTPNMLNYFGRPYEGIAVFDGKKITVTGTLWNGNKIIDERLITPDGSMLFR
VTINS

RPS9-SmBiT

MARSWVCRKTYVTPRRPFEKSRLDQELKLIGEYGLRNKREVWRVKFTLAKIRKAARELL
TLDKKDPRRLFEGNALLRRLVRIGVLDEGKMKLDYILGLKIEDFLERRLQTQVFKLGLA
KSIHHARVLIRQRHIRVRKQVVNIPSFIVRLDSQKHIDFSLRSPYGGGRPGRVKRKNAK
KGQGGAGNSGSSGGGGSGGGGSSGVTGYRLFEEIL

RPS3-LgBiT

MAVQISKKRKFVADGIFKAELNEFLTRELAEDGYSGVEVRVTPTRTEIIILATRTQNVL
GEKGRRIRELTAVVQKRFGFPEGSVELYAEKVATRGLCAIAQAESLRYKLLGGLAVRRA

CYGVLRFIMESGAKGCEVVVSGKLRGQRAKSMKFVDGLMIHSGDPVNYYVDTAVRHVLL
RQGVLGIKVKIMLPWDPTGKIGPKKPLPDHVSIVEPKDEILPTTPISEQKGGKPEPPAM
PQPVPTAGNSGSSGGGGSGGGGSSGVFTLEDFVGDWEQTAAYNLDQVLEQGGVSSLLQN
LAVSVTPIQRIVRSGENALKIDIHVIIPYEGLSADQMAQIEEVFKVVYPVDDHHFKVIL
PYGTLVIDGVTPNMLNYFGRPYEGIAVFDGKKITVTGTLWNGNKIIDERLITPDGSMLF
RVTINS

RPS3-SmBiT
MAVQISKKRKFVADGIFKAELNEFLTRELAEDGYSGVEVRVTPTRTEIIILATRTQNVL
GEKGRRIRELTAVVQKRFGFPEGSVELYAEKVATRGLCAIAQAESLRYKLLGGLAVRRA
CYGVLRFIMESGAKGCEVVVSGKLRGQRAKSMKFVDGLMIHSGDPVNYYVDTAVRHVLL
RQGVLGIKVKIMLPWDPTGKIGPKKPLPDHVSIVEPKDEILPTTPISEQKGGKPEPPAM
PQPVPTAGNSGSSGGGGSGGGGSSGVTGYRLFEEIL

RPS2-LgBiT
MGNRGGFRGGFGSGIRGRGRGRGRGRGRGRGARGGKAEDKEWMPVTKLGRLVKDMKIKS
LEEIYLFSLPIKESEIIDFFLGASLKDEVLKIMPVQKQTRAGQRTRFKAFVAIGDYNGH
VGLGVKCSKEVATAIRGAIILAKLSIVPVRRGYWGNKIGKPHTVPCKVTGRCGSVLVRL
IPAPRGTGIVSAPVPKKLLMMAGIDDCYTSARGCTATLGNFAKATFDAISKTYSYLTPD
LWKETVFTKSPYQEFTDHLVKTHTGNSGSSGGGGSGGGGSSGVFTLEDFVGDWEQTAAY
NLDQVLEQGGVSSLLQNLAVSVTPIQRIVRSGENALKIDIHVIIPYEGLSADQMAQIEE
VFKVVYPVDDHHFKVILPYGTLVIDGVTPNMLNYFGRPYEGIAVFDGKKITVTGTLWNG
NKIIDERLITPDGSMLFRVTINS

RPS2-SmBiT
MGNRGGFRGGFGSGIRGRGRGRGRGRGRGRGARGGKAEDKEWMPVTKLGRLVKDMKIKS
LEEIYLFSLPIKESEIIDFFLGASLKDEVLKIMPVQKQTRAGQRTRFKAFVAIGDYNGH
VGLGVKCSKEVATAIRGAIILAKLSIVPVRRGYWGNKIGKPHTVPCKVTGRCGSVLVRL
IPAPRGTGIVSAPVPKKLLMMAGIDDCYTSARGCTATLGNFAKATFDAISKTYSYLTPD
LWKETVFTKSPYQEFTDHLVKTHTGNSGSSGGGGSGGGGSSGVTGYRLFEEIL

LgBiT-RACK1
MVFTLEDFVGDWEQTAAYNLDQVLEQGGVSSLLQNLAVSVTLIQRIVRSGENALKIDIH
VIIPYEGLSADQMAQIEEVFKVVYPVDDHHFKVILPYGTLVIDGVTPNMLNYFGRPYEG
IAVFDGKKITVTGTLWNGNKIIDERLITPDGSMLFRVTINSGSSGGGGSGGGGSSGGAQ
KLTMTEQMTLRGTLKGHNGWVTQIATTPQFPDMILSASRDKTIIMWKLTRDETNYGIPQ
RALRGHSHFVSDVVISSDGQFALSGSWDGTLRLWDLTTGTTTRRFVGHTKDVLSVAFSS
DNRQIVSGSRDKTIKLWNTLGVCKYTVQDESHSEWVSCVRFSPNSSNPIIVSCGWDKLV
KVWNLANCKLKTNHIGHTGYLNTVTVSPDGSLCASGGKDGQAMLWDLNEGKHLYTLDGG
DIINALCFSPNRYWLCAATGPSIKIWDLEGKIIVDELKQEVISTSSKAEPPQCTSLAWS
ADGQTLFAGYTDNLVRVWQVTIGTR

SmBiT-RACK1

MVTGYRLFEEILGSSGGGGSGGGGSSGGAQKLTMTEQMTLRGTLKGHNGWVTQIATTPQ
FPDMILSASRDKTIIMWKLTRDETNYGIPQRALRGHSHFVSDVVISSDGQFALSGSWDG
TLRLWDLTTGTTTRRFVGHTKDVLSVAFSSDNRQIVSGSRDKTIKLWNTLGVCKYTVQD
ESHSEWVSCVRFSPNSSNPIIVSCGWDKLVKVWNLANCKLKTNHIGHTGYLNTVTVSPD
GSLCASGGKDGQAMLWDLNEGKHLYTLDGGDIINALCFSPNRYWLCAATGPSIKIWDLE
GKIIVDELKQEVISTSSKAEPPQCTSLAWSADGQTLFAGYTDNLVRVWQVTIGTR
